# Recovery of equilibrium free energy from non-equilibrium thermodynamics with mechanosensitive ion channels in *E. coli*

**DOI:** 10.1101/229088

**Authors:** Uğur Çetiner, Oren Raz, Sergei Sukharev, Christopher Jarzynski

**Affiliations:** Institute for Physical Science and Technology; Maryland Biophysics Program; Department of Biology; Department of Chemistry and Biochemistry; Department of Physics, University of Maryland, College Park, MD 20742; Department of Physics of Complex Systems, Weizmann Institute of Science, Rehovot 7610001, Israel.

## Abstract

Bacterial mechanosensitive channels are major players in cells’ ability to cope with hypo-osmotic stress. Excess turgor pressure due to fast water influx is reduced as the channels, triggered by membrane tension, open and release osmolytes. However, *in vitro* measurements of the free energy difference between the open and closed states of ion channels are challenging due to hysteresis effects and inactivation. Exploiting recent developments in statistical physics, we present a general formalism to extract the free energy difference between the closed and open states of mechanosensitive ion channels from non-equilibrium work distributions associated with the channels’ gating recorded in native patches under ramp stimulation protocols. We show that the work distributions obtained from the gating of MscS channels in *E. coli* membrane satisfy the strong symmetry relations predicted by the fluctuation theorems and recover the equilibrium free energy difference between the closed and open states of the channel within 1 k_B_ T of its best estimate obtained from an independent experiment.

## Introduction

The biological membrane is the fingerprint of living systems. The separation of cellular contents from the rest of the environment was the essential step for life to emerge at the expense of an osmotic stress (Haldane 1929; Oparin and Morgulis 1953; Silhavy et al. 2010), thus organisms must carefully manage the amount of internal water and osmolytes in response to variations in osmotic pressure as they change their environment in order to meet basic needs such as food and escaping from predators.

The process of converting mechanical stresses on the cell membrane into electrochemical signals is believed to be one of the oldest physiological responses of living systems to the osmotic challenges, dating back to 3.8 year billion years ago, and is well characterized in bacteria (Wood 1999; Martinac and Kloda 2003; Martinac et al. 2008; Kung et al. 2010; Wood 2015). When bacteria are exposed to hypo-osmotic stress, e.g. during a rain storm, the turgor pressure builds to dangerous levels in a few milliseconds due to high membrane water permeability (Wood 1999; Boer et al. 2011; Çetiner et al. 2017). One of the main mechanisms to evade mechanical rupture due to extreme internal turgor pressure is the activation of mechanosensitive ion channels (MS) in the membrane. These reduce excess turgor by releasing internal osmolytes and water (Fig. 1A). In bacteria, the bulk release of ions and other osmolytes is mainly mediated by two families of mechanosensitive channels: MscS and MscL. The 3-nS MscL family channels gate at near lytic tension and therefore serve as a “last back up” mechanism against extreme osmotic down shocks by forming large non-selective pathways in the membrane (Sukharev et al. 1994; Sukharev et al. 1999; Chiang et al. 2004). The MscS channels, on the other hand, have a conductance of 1 nS, require less tension to open and come with great diversity in structure and functionality (Malcolm and Maurer 2012; Cox et al. 2015).

**Figure 1.**
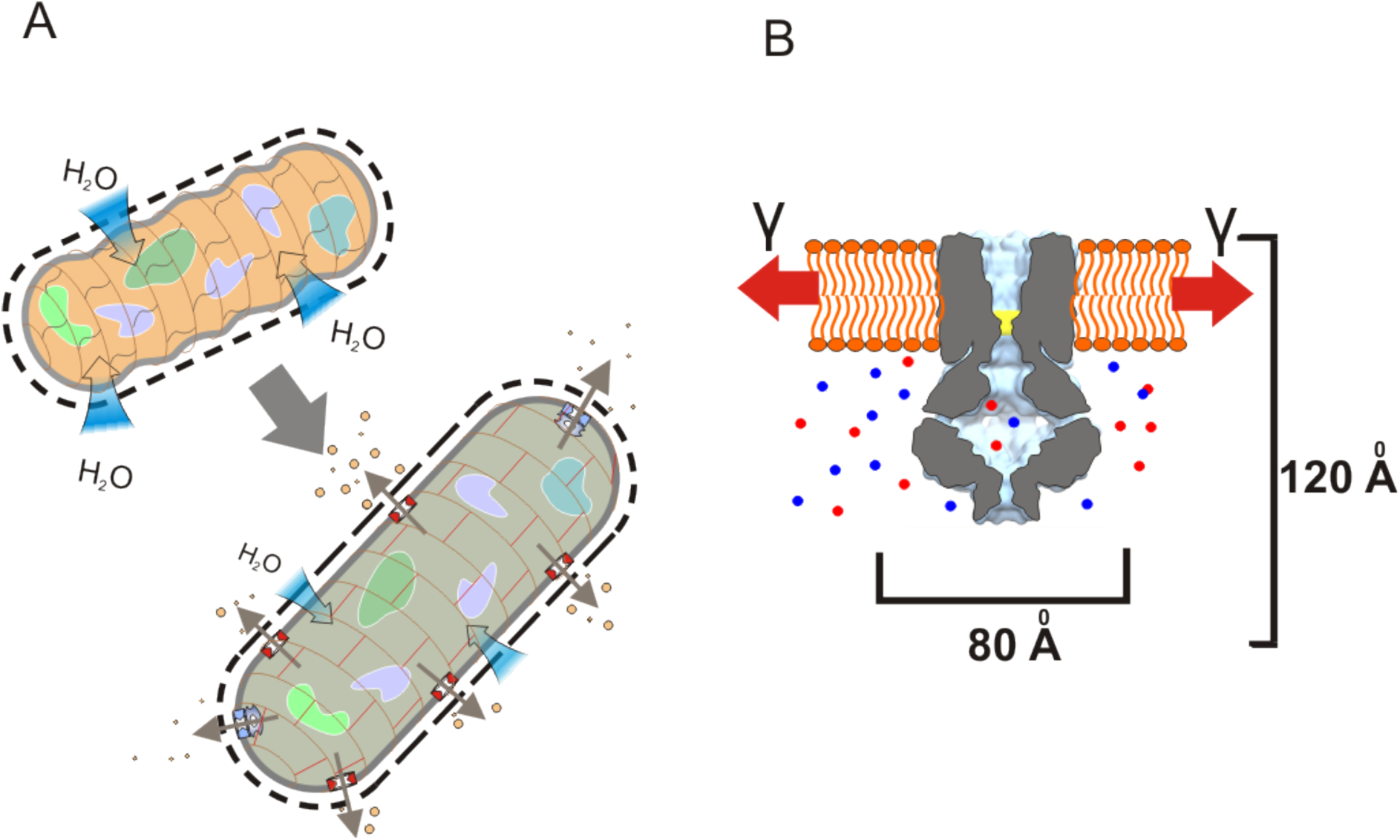
**(A)** Bacterial response to osmotic downshock. Water influx is accompanied by elastic deformation of the cell wall and stretching the cytoplasmic (inner) membrane. Mechanosensitive channels open and release small osmolytes together with water. When tension and volume return to normal, channels close. **(B)** Cartoon of MscS channel in the closed confirmation. As tension increases, the free energy difference between the open and closed state decreases thus in the presence of membrane tension, the open state becomes energetically more favorable.

MscS deserves special attention because it senses not only the magnitude of tension but also the rate of tension application and responds differently to abrupt pulses and slow ramps (Akitake et al. 2005; Belyy et al. 2010). It appears to be optimized for tension stimuli ranging in length from tens of seconds, where it adapts and completely equilibrates between the functional states (Cetiner and Sukharev 2017), to milliseconds, where it acts in a highly non-equilibrium regime with a drastic disparity between fast opening and slow closing kinetics. As will be discussed below, this asymmetric (non-equilibrium) behavior of MscS can be a part of its native mechanism as it well suits its functional role of low-threshold release valve in a small bacterial cell where water and osmolyte exchange rates are largely dictated by membrane water permeability and the surface area-to-volume ratio.

Recent findings have suggested that in order to survive under extreme hypo-osmotic conditions, the rate of turgor release via mechanosensitive ion channels should keep up with the rate of turgor generation caused by the water influx (Bialecka-Fornal et al. 2015; Buda et al. 2016; Çetiner et al. 2017). According to this mechanistic picture, the osmotic water permeability of the cell envelope and the magnitude of the shock determine the rate of water influx whereas parameters such as channel density, conductance, and the free energy difference between the closed and open confirmations (ΔF) govern the rate of osmolyte release.

In order to develop a quantitative understanding of survival and fitness in microbial world under extreme osmotic stress, it is crucial to obtain accurate information on all the relevant parameters of the MS channels. The free energy difference ΔF, an indispensable part of the electrophysiological characterization of the channels, is of particular interest to experimentalists. The most common way to measure ΔF is to ramp the membrane tension linearly in time until all the channels are open, while measuring the conductivity between the two sides of the membrane. These traces of conductivity vs. tension – the dose response curve – correspond (when properly normalized) to the probability of finding the channel in “open” or “closed” states. This probability is fitted to a two-state Boltzmann distribution function, and the free energy difference is extracted from the distribution (Sukharev et al. 1999; Chiang et al. 2004; Nakayama et al. 2013; Petrov et al. 2013). A crucial assumption in this method is that the distribution of states is accurately described by the Boltzmann distribution. Whereas this is a perfectly valid assumption for a system in thermal equilibrium, it might not hold for systems out of equilibrium. Indeed, if the protocol is delivered in a time symmetric manner such that the membrane tension is increased and decreased with the same rate as in a triangular ramp protocol, the current follows different paths on the ascending and descending legs of the dose response curve and displays a clear hysteresis loop – the fingerprint of nonequilibrium processes. The choice of equilibrium-based formalism to extract the free energy difference for a non-equilibrium process may contribute to the variability in ΔF in the literature, ranging from 5 *k_B_T* to 28 *k_B_T* for the same MscS channel (Kloda and Martinac 2002; Wang et al. 2008; Belyy, Kamaraju, et al. 2010).

In many cases, by delivering the stimulus slowly it is possible to obtain the free energy difference from the ‘’reversible” work, W_*rev*_ ≅ ΔF. However, there are channels such as *E. coli* MscS and PaMscS-1 that undergo inactivation and become non-conductive and insensitive to tension as they are pulled slowly. This phenomenon, which is called “the Dashpot Mechanism”, prevents data acquisition at a slow enough rate to assume thermal equilibrium (Akitake et al. 2005; Çetiner et al. 2017).

In this article, we present alternative and more reliable ways to obtain the free energy difference between the open and closed state of MscS by making use of recent developments in statistical physics. In 1997, Jarzynski showed that it is possible to obtain the free energy difference between two equilibrium states (A, B) from the non-equilibrium distribution of the work performed on the system during a thermodynamic process connecting A to B (Jarzynski 1997):

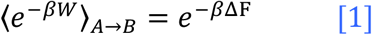

Here the angular brackets represent the ensemble average taken over many realizations of the same switching protocol starting from an equilibrium state (A) in contact with a heat reservoir at temperature T. *β* is defined as 1/*k_B_T* where *k_B_* is the Boltzmann constant. In 1999, Crooks proved that the work distributions associated with the thermodynamic process of switching the system from A to B and with the corresponding time-reversed protocol of switching the system from B to A (where the system starts in the corresponding equilibrium state, A or B) satisfy the following symmetry relation (Crooks 1999):

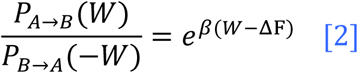

These fluctuation theorems are robust regardless of how microscopic dynamics are modeled (Hummer and Szabo 2001; Hatano and Sasa 2001; Sun 2003; Evans 2003; Oberhofer et al. 2005; Imparato and Peliti 2005; Seifert 2005), and they have been experimentally verified in various systems (Liphardt et al. 2002; Wang et al. 2002; Douarche et al. 2005; Collin et al. 2005; Blickle et al. 2006; Harris et al. 2007; Saira et al. 2012; An et al. 2014).

Using these theoretical results, we present a general framework that enables one to extract the free energy difference between the open and closed state of ion channels in the patch clamp experiments from the non-equilibrium generalized work distributions associated with the gating of mechanosensitive ion channels. Having several channels in the membrane provides the advantage of obtaining many data points in a single realization of the experimental protocol. Thus, the statistics from which the work distributions can be extracted enables fast convergence for eqns [1] & [2] (Gore et al. 2003; Jarzynski 2006). Moreover, a recent study, which modeled MscS gating as a discrete state continuous-time Markov chain (Cetiner and Sukharev 2017), enabled us to validate these fluctuation theorems using mechanosensitive ion channels in a patch clamp setup.

## Experimental and Theoretical Setup

The patch clamp technique was developed by Erwin Neher and Bert Sakmann in the late 70s and early 80s (Neher and Sakmann 1976; Hamill et al. 1981; Sakmann and Neher 1984). It enables researchers to characterize the electrophysiological properties of ion channels by clamping a piece of a membrane as a giga-ohm seal in a polished glass micropipette (Fig. 2A). The high resistance of the seal provides an electrical isolation across the membrane. However, conducting pathways can be generated by activation of mechanosensitive ion channels in response to applied tension. This activation can be monitored with pico-amp precision. Observations of discrete currents passing through individual channels made patch-clamp essentially the very first single-molecule technique.

**Figure 2.**
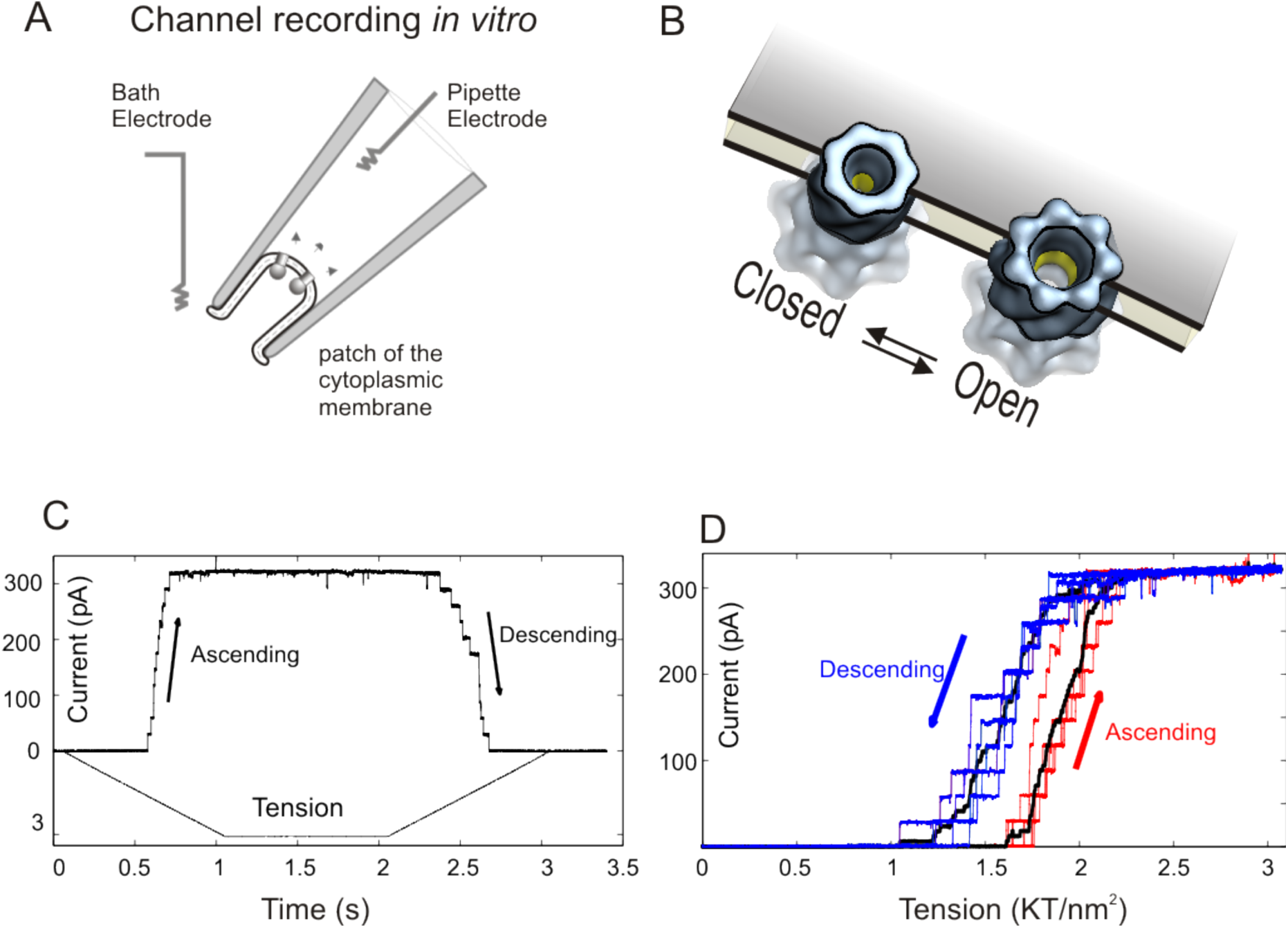
**(A)** Patch-clamp applied to giant bacterial spheroplasts has been the most informative way of obtaining functional characteristics of bacterial mechanosensitive channels (Martinac et al. 1987). A micron-size glass pipette holds a cup-shaped patch separating the inside of the pipette from the bath. Application of suction to the pipette stretches the curved patch membrane according to the law of Laplace and generated tension activates the channels. The two electrodes inside and outside the pipette measure the current reflecting the opening and closing transitions in individual channels. **(B)** A blow-up of a fragment of the patch membrane showing two channels, one on the closed and one in the open state. The open channel has a larger cross-sectional area in the plane of the membrane. As membrane tension increases, the free energy difference between the open and closed state decreases thus in the presence of membrane tension, the open state becomes energetically more favorable. **(C)** An example of typical trace containing 11 MscS channels and the tension stimulus shown below. On ascending leg, membrane tension was increased linearly, ***γ*** = ***rt*** at a rate ***r*** = ***3 k_B_T/nm^2^s^−1^*** until all the channels in the membrane opened, manifested as the saturation of the current. After 1s of equilibration at ***3 k_B_T/nm^2^*** the tension was decreased back to 0 ***k_B_T/nm^2^*** at the same constant rate, r. The descending leg depicts the closure events of the channels. **(D)** Five representative traces from the same patch were plotted as function of tension in order to emphasize that the single channel events are stochastic and the system displays hysteresis as demonstrated by the averaged traces (black curves).

In our experiments, the system of interest is the collections of mechanosensitive ion channels naturally embedded in the *E. coli’s* membrane. The micropipette with the clamped membrane is immersed into the bath solution at room temperature, which serves as a thermal reservoir. The “work parameter” (see definition below) is the membrane tension, γ and the conjugate variable to work parameter is the lateral protein area expansion, *A* (Fig. 2B). The application of suction changes the pressure between the two sides of the membrane hence varies its tension, which allows us to perform work on the system and lower the free energy difference between the open and closed state. Thus, in the presence of external tension on the membrane, states with larger area become favorable (Sukharev et al. 1999; Kamaraju et al. 2010).

A typical experimental protocol starts with a membrane without any tension in which all ion channels are in the closed state and the conductance is negligible. Then, a linear increase in the membrane tension to a value of about 3 *k_B_T/nm^2^* in 1s is applied. This tension is kept constant for another second to let the system relax to an equilibrium state corresponding to the final value of the work parameter. Once the system has reached its equilibrium distribution, the tension is decreased back to 0 *k_B_T/nm^2^* at the same rate. Fig. 2C displays a characteristic response of the current to the tension protocols described above: each step of current increase (decrease) represents a single channel opening (closing) event. In the specific example of Fig. 2C there are 11 mechanosensitive channels which are open at the final tension equilibrium state. Note that the longer the membrane is exposed to high tension values such as 3 *k_B_T/nm^2^*, the greater the chance that it ruptures. Therefore, 1s holding period is a safe time scale not only to reach the equilibrium (assuming that the system’s relaxation time at 3 *k_B_T/nm^2^* is a few milliseconds) but also to obtain a few realizations before the seal is lost (see SI Fig. 1). Fig. 2D shows five representative current traces of the same protocol, applied on the same membrane. We emphasize that: (i) Even though the same protocol is employed, the channel gating is fluctuating. We will show that these fluctuations satisfy strong symmetry relations. (ii) As indicated by the trajectories average currents (the black curves Fig. 2D), the system displays clear hysteresis, which is the hallmark of non-equilibrium processes. Therefore, assuming that the distribution of open and close states follows the Boltzmann distribution at all times is not a valid assumption, and nonequilibrium fluctuation theorems would be more suitable to extract the free energy difference between the open and the closed state of the channel.

We model MscS as a two-state system (“open” and “closed”), and introduce a state variable, σ, which is 0 for a closed channel and 1 for an open one. Let *ε_closed_* and *ε_open_* denote the energies of the closed and open states. In the presence of tension *γ*, the energy of the system can be written as (Ursell 2009):

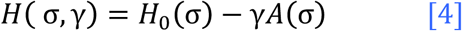

where the term *H*_0_(σ) = (1 — σ) *ε_closed_* + σ*ε_open_* represent the energy of the system due to the state of the ion channel alone, and the additional term *γA*(σ) = *γ*σΔ*A* represents the energy difference between open and close channels in the presence of tension, where Δ*A* is the area difference between the closed and open state of the channel.

For a single realization of the experimental protocol of duration *τ*, the work is defined as the energy difference associated with the changes in the external tension:

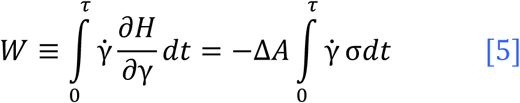

This work takes on different values in different realizations, namely it fluctuates from one realization to another. We should note that eqn. [5] is the work definition used in the context of non-equilibrium thermodynamics and it differs from the classical textbook definition of mechanical work. (See (Jarzynski 2007; Horowitz and Jarzynski 2007) for a detailed discussion). Writing eqn. [5] in discrete time steps is convenient for our purpose, since the experimental acquisition yields digitized data in discrete steps, whose size depends on the sampling frequency. Considering that the tension (*γ*) is increased linearly from zero to it is final value *γ_Final_* in M small enough time steps, a dynamical path for a single channel might be described by the following sequences:

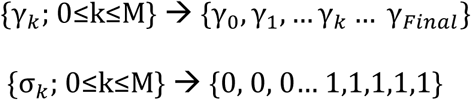

The work performed on the system, eqn [5], can be rewritten in discrete time for a single realization as (Ritort et al. 2002):

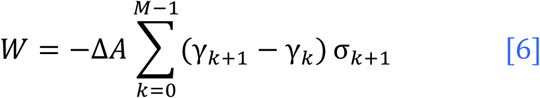

A typical realization, for a cyclic protocol, is depicted in Fig. 3 where the red curve represents the opening event and the blue curve is the closing event. For the sake of simplicity, let us assume Δ*A* = 1 *nm^2^*. In this case, for the ramp increase in the tension, eqn [5] implies that the work done is minus the area under the σ vs *γ* curve. Specifically, the work performed in order to open the channel, *W_C→0_*, is equal to minus the area under the red curve, which is denoted by — *W_0_*, and the work performed during the closure *W_O→C_* equals to the area under the blue curve, *X* + *W_O_*, where X is the area bounded by the two curves (See Fig. 3). Therefore, the total dissipation during the thermodynamic cycle is *W_Diss_* = *W_C→0_* + *W_0→C_* = *X* > 0. In other words, the average work done in order to open the channel is greater than the average work performed as the channel closes. However, in some rare realizations the thermal noise might act on the channel “constructively” and the channel might gate at lower work values than the free energy difference, *W* < ΔF, violating the second of thermodynamics transiently.

**Figure 3.**
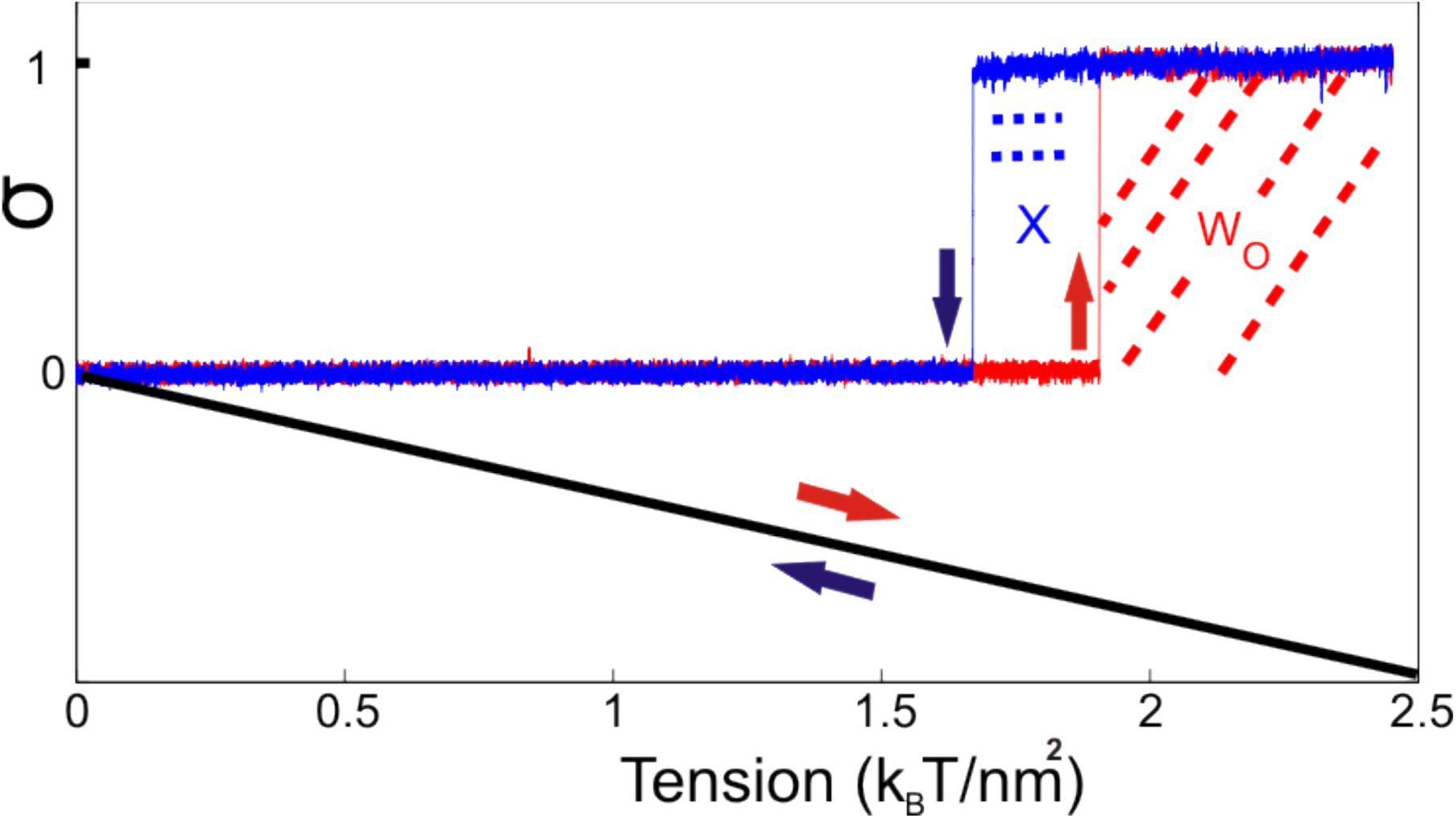
The gating (red) and closure (blue) of a single channel in response to linear increase and decrease of the membrane tension with the same rate. For the sake of illustration, assume **Δ*A*** =1 nm^2^. In this case 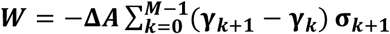 gives the work performed during the gating, ***W_C→0_***, as minus the area under the red curve (—***W_0_***) and the work performed during the closure ***W_0→c_*,** as the area under the blue curve (***W_O_* + *X***). The total dissipation for this cyclic process is ***W_Diss_*** + ***W_C→O_*** + ***W_O→C_*** = ***X* > 0** which holds on average for small systems. For some rare realizations, it is possible to receive more work than what is put in giving ***W_Diss_*** < **0**. Imagine a realization where the colors and the direction of the arrows are swapped and the red curve now represents the closure and blue curve depicts the gating, in this case:

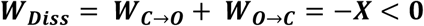

## Results

An edge detection program was used to identify single channel events in all the traces (see SI Fig. 2). The corresponding work distributions associated with the channel’s opening at a given tension protocol, *P_c→0_*(*W*), and the closing work distribution at the time reversed protocol, *P_0→c_*(—*W*), were obtained from two independent experiments, and are depicted in Fig 4. The work values were binned into 13 equally spaced intervals. Note that according to eqn [2], at the work value W_0_ for which *P_c→0_*(*W*_0_) = *P_0→c_*(—*W*_0_), it holds that W_0_ = ΔF. In other words, the work distributions of the forward and backward protocols cross at W = ΔF. This observation, although quite convenient, does not provide an optimal method to extract ΔF since it only uses local behavior of the work distributions around the crossing point, whose exact location depends on the choice of the bin size. Instead, the plot of log(*P_c→0_* (*W*)/*P*_0→c_(—*W*)) as a function of W/*k_B_T* provides a better estimate as it relies on the whole distribution. The points on the semi-log plot follow a straight line with slope 1 and intercept the work axis at ΔF. Based on the plotted measurements, the free energy difference is estimated to be −15.3 *k_B_T* from the crossing point in Fig 4 while the straight line, in supplementary Fig. 3A intercept work axis at −14.7 *k_B_T* with the slope of 0.98.

**Figure 4.**
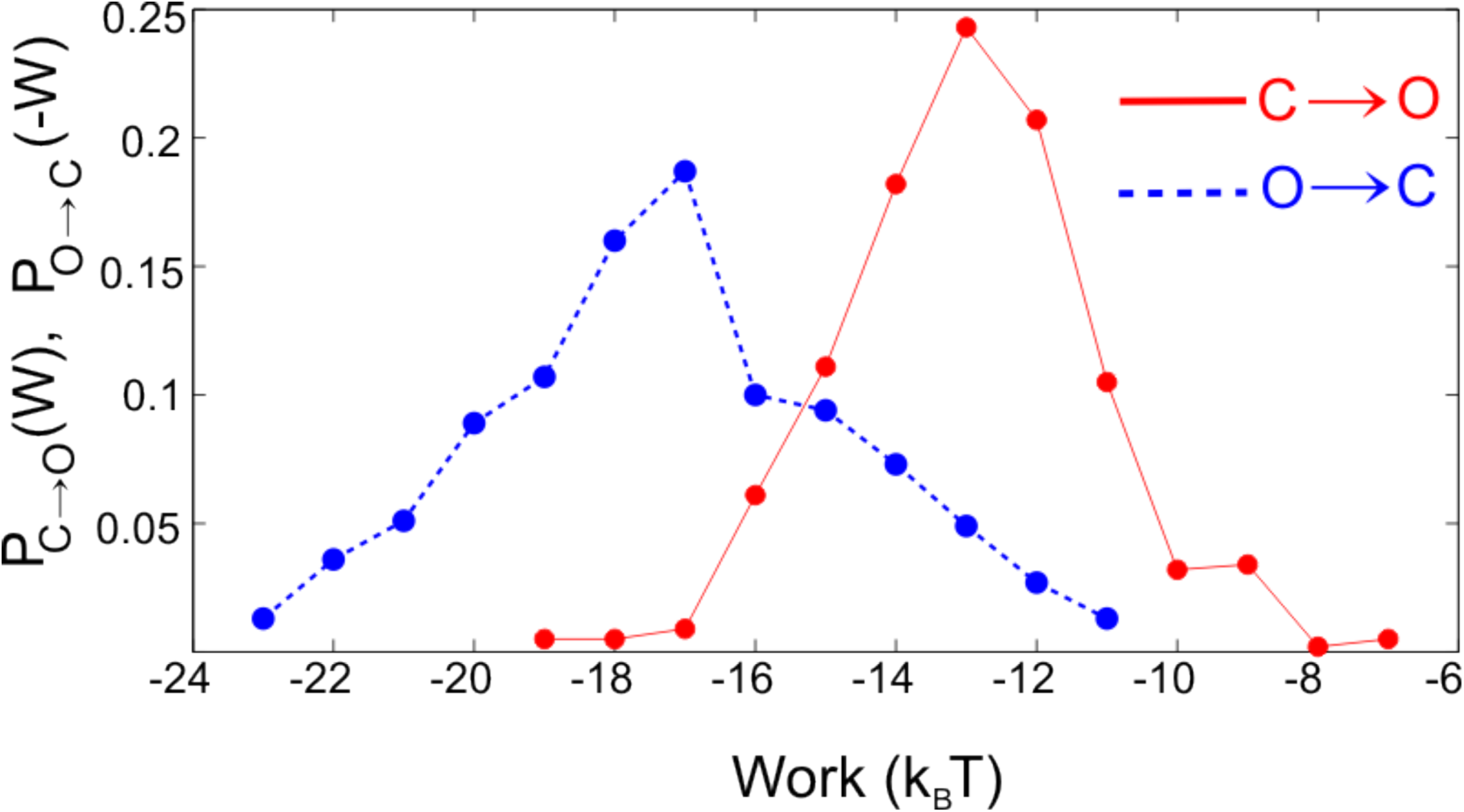
Histograms of the work distrubutions at **3 *k_B_T/nm^2^s*^−1^**. N=440 for the opening (***C* → *O***) and N=449 for the closing (***O*** → ***C***). The protocol was carried time symmetrically. The work distributions cross at −15.3 ***k_B_T***. The individual work values were obtained from the following definition: 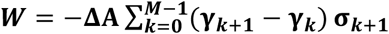.

It should be noted that even though the plot of log ratio of the work distributions as a function of work exploits the full distributions and offers an improvement, the overall probabilities still depend on the binning size. Moreover, this method is only useful on the condition that there is a significant overlap between the work distributions, therefore for highly dissipative processes, the overlap would be quite poor and alternative statistical techniques such as Bennett’s acceptance ratio method (see Supplementary Information, Fig. 3B) can be used to recover ΔF (Bennett 1976; Crooks 2000; Shirts et al. 2003).

Jarzynski equality (Eq. [1]) for the closed → open transition, 〈*e*^−*βw*^〉_C→0_ = *e*^−*β*ΔF^, and for the open → closed transition, 〈*e*^−*βw*^〉_0→*C*_ = *e*^−*β*ΔF^, provide another estimation for ΔF. These give −14.9 ± 0.4 *k_B_T* and 14.1 ±0.4 *k_B_T*, respectively. Thus, the average, −14.5 ± 0.3 *k_B_T* is the ΔF predicted by Jarzynski equality. The overlapping distribution method (Bennett 1976; Frenkel et al. 1997) is also in good agreement with other estimators and reports ΔF to be −14.7 *k_B_T* (See Supplementary Information, Fig. 4). Finally, the Linear Response Theorem (LR) recovers the free energy from the average work done during the opening or from the average work received during the closing and the corresponding standard deviations of the work distributions as follows (Hermans 1991):

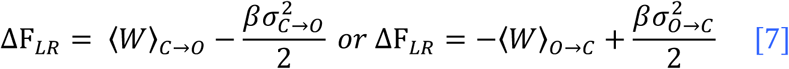

which results in ΔF_*LR*_ = —14.7 ±0.2 *k_B_T* and −13.8 ±0.3 *k_B_T*, respectively. The summary of the results produced by different estimators is listed in Table 1 with standard deviations in units of k_B_T. Uncertainties were obtained using the bootstrap method (Efron 1979).

**Table 1.** **Summary of Results. 〈W_Diss_〉= 〈W〉_c→0_ + 〈W〉_0→C_,** is the average dissipation during the cyclic process (C→O→C), The linear response prediction and Jarzynski estimations are the averages of the free energy differences obtained for the opening and closing current trajectories. ΔF from the intersection of the work distributions was predicted to be −15.3 ***k_B_T***. A better estimate is achieved by using the semi-logarithmic plot of Crooks’ Fluctuation Theorem which gives ΔF = −14.7 ***k_B_T*** (See SI Fig. 3A], Overlapping distribution method is in line with other estimators and predicts ΔF = −14.7 ***k_B_T***. The Bennett’s acceptance ratio method, which is our best guess for the free energy, reveals it to be −15.0 ± **0. 5 *k_B_T***. (See the Supplementary Information, Fig. 3B]

| $\langle W \rangle_{C \rightarrow O}$ | $\sigma_{C \rightarrow O}$ | $\langle W \rangle_{O \rightarrow C}$ | $\sigma_{O \rightarrow C}$ | $\langle W_{Diss} \rangle$ | $\Delta F$<br>Linear<br>Response | $\Delta F$<br>Jarzynski | $\Delta F$<br>Crooks | $\Delta F$<br>Bennett's | $\Delta F$<br>Crossing | $\Delta F$<br>Overlapping |
| --- | --- | --- | --- | --- | --- | --- | --- | --- | --- | --- |
| -13.0 | 1.82 | 17.2 | 2.60 | 4.2<br>$\pm 0.2$ | -14.2<br>$\pm 0.2$ | -14.5<br>$\pm 0.3$ | -14.7<br>$\pm 0.2$ | -15.0<br>$\pm 0.5$ | -15.3<br>$\pm 0.6$ | -14.7<br>$\pm 0.2$ |

It is important to note that the free energy estimations in Table 1 refer to equilibrium free energy differences between two equilibrium states associated with the initial and final value of the work parameter (system + driving force) (Douarche et al. 2005). For a typical experimental protocol in this paper, Δ*F* ≡ F (*γ_τ_* = 3 k_B_T/nm^2^)- F (*γ*_0_= 0 k_B_T/nm^2^), whereas Δ*F*_0_ represents the free energy difference between the closed and open state of the channel in the absence of any tension. Therefore, even though Δ*F* depends on the end points of the work parameter, Δ*F*_0_ should be independent of the protocol and can be recovered by adding the corresponding boundary term to Δ*F*:

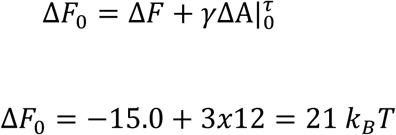

Here the final value of the tension, *γ*(*τ*) = 3 *k_B_T/nm^2^* and ΔA was taken to be 12 *nm*^2^ (Boer et al. 2011; Nakayama et al. 2013). With totally different techniques, Δ*F*_0_ was determined to be 22 ± 1 *k_B_T* (Cetiner and Sukharev 2017).

## Discussion

We extracted the free energy difference between the closed and open state of a mechanosensitive ion channel (MscS) in *E. coli’s* membrane from non-equilibrium work measurements performed in the patch clamp setup where the work parameter is the membrane tension, *γ*, and corresponding conjugate variable is the protein’s lateral area expansion, Δ*A*. Various estimators are all in good agreement with each other and recover the free energy within *k_B_T* of its best estimate obtained from an independent experiment. We considered the Bennett’s acceptance ratio method as our best estimate for the free energy since it uses both of the work distributions associated with *C* → *O* and *O* → *C* transitions and it does not rely on binned histograms between the distributions.

The general framework presented in this paper is not limited to MscS but is valid for all mechanosensitive ion channels that can be modeled as two-state systems. This condition can usually be met by a fast delivery of the membrane tension at the expense of irreversibility and dissipation, which does not pose a problem since nonequilibrium fluctuation theorems can still recover the free energy for the processes where *W_Diss_* ≤ 100 k_B_T (Collin et al. 2005). It is also possible to extend eqn [6] for the voltage sensitive channels where the work parameter is the electric field across the membrane and the conjugate variable is the gating charge (q). In this case, the open state becomes favorable in the presence of external voltage across the membrane, as the movements of charged residues in the electric field, qV, lowers the free energy difference between the open and closed configurations.

It is also worth mentioning that even though the work parameter (the external tension) changes as a function of time according to a predefined protocol, in principle it should not respond to the state of the system, namely it should not fluctuate and should be identical in each realization of the protocol. This criterion can only be satisfied in simulation or theory. This problem is not unique to our setup. It appears as well in single molecule pulling experiments (Manosas and Ritort 2005; Wen et al. 2007; Manosas et al. 2007), where the work parameter is the molecular extension of the system or the force calibrated from the displacement of the bead in a trap. Neither of these parameters is exactly the same for each repetition of the experiment due to different response of the pulled molecule, thus, the assumption that these parameters are controlled is an approximation (Ritort 2007). By the same token, the work parameter in the patch clamp experiment is not immune to fluctuations. Therefore, in any experimental setup, extra care should be taken to ensure that the work parameter is properly chosen such that the fluctuations are negligible. In Supplementary Fig. 5, we compared the work distributions and free energy difference obtained from two separate membranes having different number of MscS channels. A good reproducibility has been observed among different membranes under the same protocol. Moreover, we tried a different protocol where the membrane tension was increased to 3.6 *k_B_T/nm^2^* in 250 milliseconds giving a different Δ*F*. As a control experiment, regardless of the final value of the work parameter or the rate, Δ*F*_0_ should always be retrieved from 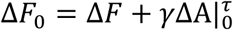 since the boundary term compensates for the difference in Δ*F* (see Supplementary Fig. 6 and Table-2). It is also important to note that the framework developed in this paper can still recover Δ*F*_0_ for the channels where Δ*A* is not known in advance. If we repeat the measurements for different values of *γ_τ_*, the change in Δ*F* with respect to *γ_τ_* reveals Δ*A*, which can then be used to retrieve Δ*F*_0_ as shown above or in SI.

It is an experimental advantage to have as many channels as possible, since each single channel event yields an independent work value for every realization of the experimental protocol. Yet, many channels come with two inevitable complications inherent in all patch clamp experiments. (i) To use the proposed method single channel events must be resolved. This becomes increasingly difficult when there are many channels in the same patch. Therefore, we expressed MscS in a tightly regulated PBAD expression system in order to control channel expression (ii) If a channel changes its state more than once, the path it follows, {σ_*k*_; 0≤k≤M}, cannot be determined precisely for the rest of the protocol, since the single channel events are indistinguishable in the patch clamp setup. Fig. 5A demonstrates an example of the second complication where the work associated with the transient gating of a channel in the beginning of the experiment might belong to any of these three channels. Similarly, in Fig. 5B, one of the channels at the end of the protocol briefly changes its state from open to closed as shown by black and red arrows. If such events, although not very common, happen in the beginning or at the end of the protocol, the error introduced will not be significant as the area between the red and black arrows is negligible compared with the typical area that gives the work values. Despite this unavoidable complication, the patch clamp technique is still quite advantageous and offers fast convergence as enough statistics for the work distributions can be collected efficiently.

**Figure 5.**
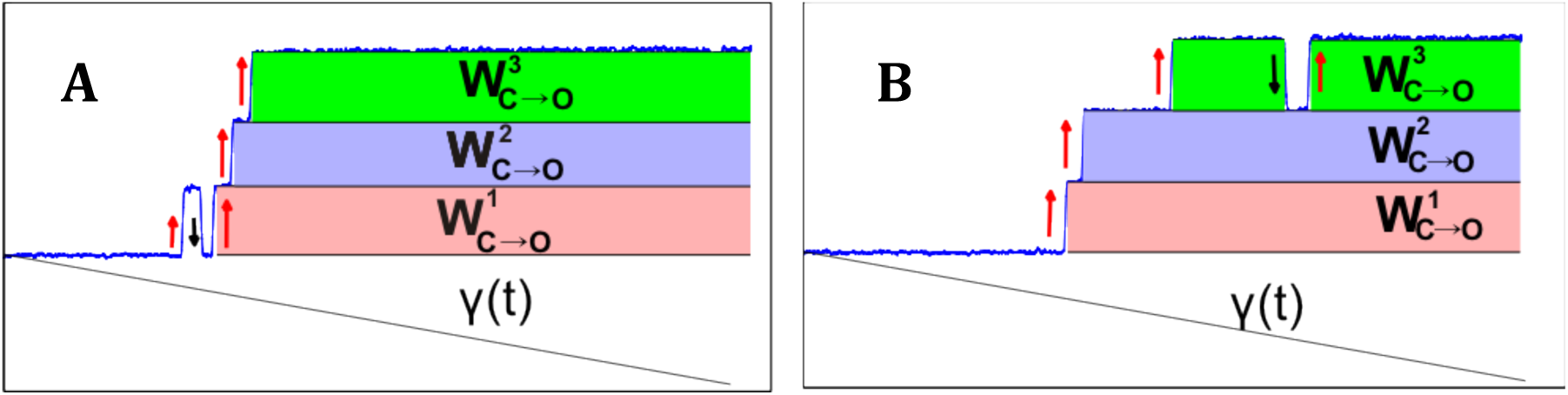
An inevitable complication due to indistinguishability of single channel events in the patch clamp experiments. Gating of 3 MscS channels in response to linear increase in the membrane tension is depicted. For the sake of simplicity, assume that **Δ*A*** =1 nm^2^, therefore the shaded areas represent the work done during the gating. The transient opening **(A)** and closing **(B)** events denoted by the red and black arrows lead to an error that is proportional to the area between the arrows. More specifically, the work done for the one of the channels in (A) is underestimated whereas in B, overestimated by an amount corresponding the area between the red and black arrow. This error still lies within experimental uncertainty and can be neglected compared to typical work values obtained from the shaded areas.

The value of equilibrium free energy change ΔF for a particular process is two-fold. First, it provides both the limit of recoverable energy when this molecular machine is exploited under ideal reversible conditions and the measure of dissipation in a real process. Second, and more importantly, it represents the path-independent thermodynamic parameter attributable to the transition between the two specific end conformations and thus is a unique characteristic of that specific change in molecular structure. The data presented above shows that the 1 s linear ramp regime (at a rate r = 3 k_B_T/nm^2^ s^−1^) is not too far from the true equilibrium regime: the energy dissipated due to forced opening is on average about 4 *k_B_T* on top of the equilibrium ΔF measured as 21 *k_B_T*. The observed hysteresis between the opening and closing trajectories becomes stronger when the rate of stimulus application and release increases (SI Fig. 7). The question is in what regime the channel functions in vivo. Our recent kinetic studies of the light scattering signal in stopped-flow experiments indicated that in a wild-type *E. coli* cell possessing ~70 MscS and ~50 MscL channels an abrupt 900 mOsm downshift liberates nearly 15% of all internal metabolites within 30 ms with only a small viability loss (Çetiner et al. 2017). In this regime, the channels activate within 15 ms of tension onset, the time which approximates the time course of “native tension ramp” generated by osmotic swelling. This activation regime is much farther from equilibrium than any of the patch-clamp regimes presented here. We may conclude that the manifestations of even more pronounced hysteresis would be fast activation (opening) possibly by slightly elevated tension. However, if membrane tension drops to near-threshold tension in the 15-30 ms time scale then it will be followed by a relatively slow closure allowing for more complete solute equilibration between the bacterial cytoplasm and the medium even when the tension drops below the activation threshold. Therefore, the non-equilibrium gating of MscS in vivo appears to be a beneficial trait embedded in the channel mechanism.

## Material and Methods

### Preparation of giant spheroplasts and patch clamp

The giant spheroplasts of *E. coli* were prepared following the protocol described in (Martinac et al. 1987). 3 ml of the colony-derived culture was transferred into 27 ml of LB containing 0.06 mg/ml cephalexin, which selectively blocks septation. After 1.5-2 hours of shaking in the presence of cephalexin, 100-250 μm long filaments formed. Toward the end of the filamentous growth stage, induction with 0.001 % L-Arabinose was done for 10-20 mins which gave 1-15 channels per patch. The filaments were transferred into a hypertonic buffer containing 1 M sucrose and subjected to digestion by lysozyme (0.2 mg/ml) in the presence of 5 mM EDTA. As a result, filaments collapsed into spheres of 3-7 μm in diameter in 7-10 mins. The reaction was terminated by adding 20 mM Mg^2^+. Spheroplasts were separated from the rest of the reaction mixture by sedimentation through a one-step sucrose gradient.

Borosilicate glass (Drummond 2-000-100) pipets 1-1.3 *μ*m in diameter were used to form tight seals with the inner membrane. The MS channel activities were recorded via inside-out excised patch clamp method after expressing them in MJF641. The pipette solution had 200 mM KCI, 50 mM MgCI_2_, 5 mM CaCI_2_, 5 mM HEPES. The bath solution was the same as the pipette solution with 400 mM sucrose added. Both pipette and bath solution had the pH of 7.4. Traces were recorded using Clampex 10.3 software (MDS Analytical Technologies). Mechanical stimuli were delivered using a high-speed pressure clamp apparatus (HSPC-1; ALA Scientific Instruments).

### Tension Calibration

The pressure (P) was converted to the tension (*γ*) using the following relation: *γ* = (*P*/*P*_0.5_)*γ*_0.5_ assuming the radius of curvature of the patch does not change in the range of pressures where the channels were active (P> 40 mmHg) and the constant of proportionality between tension and pressure is *γ*_0.5_/*P*_0.5_ (Sukharev et al. 1999; Moe and Blount 2005; Belyy, Kamaraju, et al. 2010). The midpoint tension, *γ*_0.5_ of MscS was taken to be 7.85 mN/m (Belyy, Kamaraju, et al. 2010). *P*_0.5_ represents the pressure value at which half of the population is in the open state and was determined from the averages of 5-10 traces obtained by using 1-s triangular ramp protocols at the beginning of each experiments.

## Acknowledgments…

UC was supported by the U.S. Department of Education GAANN “Mathematics in Biology” Fellowship. OR is supported by a research grant from Mr. and Mrs. Dan Kane and the Abramson Family Center for Young Scientists. The work was supported by NIH R21AI105655 and GM107652 grants to SS. CJ acknowledges financial support from the National Science Foundation (USA), under grant DMR-1506969. UC thanks Ms. Stephanie Sansbury for cloning MscS into tightly-regulated pBAD for expression system and Dr. Andriy Anishkin for the illustrations in Fig. 1 & 2. UC is also indebted to Dr. Yigit Subaçi for the stimulating discussions.

